# Can force-plate measurement be trusted for balance diagnostics? Frequency-domain force-plate performance assessment for quiet-standing studies

**DOI:** 10.64898/2026.07.07.737003

**Authors:** Rika Sugimoto-Dimitrova, Jiajie Qiu, Neville Hogan

## Abstract

Older adults face an increased risk of falls that may have severe consequences for their well-being. Routine, accessible clinical screening may help mitigate fall risk through early detection of balance impairments. Portable force plates offer a convenient and practical solution for balance assessment in clinical settings. A new force-plate-based balance measure, the intersection-point-height, has shown particularly promising results in its ability to distinguish between healthy and impaired balance behaviors. However, the intersection-point-height measure requires measurement of shear force during standing, which exhibits magnitudes of less than 0.2% of normal forces (body weight), taxing the dynamic range of most sensor technologies. The ability of existing force plates to measure such low-magnitude shear forces observed during quiet standing is currently unknown. This study presents a force-plate performance assessment method to evaluate shear-force measurement errors and quantify the uncertainty of the intersection-point-height measure. The method was applied to test a commonly used laboratory-grade portable force plate. While the device successfully captured sagittal-plane intersection-point-height at the lowest frequencies, low signal strength prevented precise readings in the frontal plane. Thus, the tested device only marginally met the precision required for quiet-standing analysis, underscoring the critical need for systematic performance validation of portable force plates prior to clinical use. Future efforts should focus on evaluating alternative portable force plates and exploring economical design improvements to enhance shear-force measurement precision.

## 1 Introduction

Unintentional falls among older adults in the United States resulted in 41,400 deaths and 1.32 million hospitalizations in 2023, with medical costs amounting to $73 billion (Centers for Disease Control and Prevention). These numbers increased over 40% in the last 8 years due to the growing older-adult population. As the world’s population ages, there is a pressing need to address fall risk among older adults.

Fall prevention requires early detection of balance impairment. This effort would benefit from a “vital signs” measure of balance ability to be incorporated into the routine check-up performed at a doctor’s office. New and potentially informative data is available from shear force during standing: the intersection-point-height is a frequency-domain measure related to the covariance between shear force and center-of-pressure displacement during quiet standing, and can detect differences in balance behavior with age, Parkinson’s disease, and stroke (Boehm et al. 2019, Shiozawa et al. 2024a, Sreenivasan et al. 2024, Bartloff et al. 2025, Shiozawa et al. 2026). However, shear force observed during quiet standing is several orders of magnitude smaller than body weight. Can measurements of shear force be made with acceptable reliability and low-enough cost to belong in every clinician’s office?Although center-of-pressure measurement uncertainty has been extensively studied (Bobbert & Schamhardt 1990, Middleton et al. 1999, Chockalingam et al. 2002, Morasso et al. 2004, Cedrado et al. 2009, Blanchard Sanhueza 2010, Ghersi et al. 2017, List et al. 2017, Limpach et al. 2025) no study has evaluated shear-force uncertainty for force magnitudes expected while standing. Those that did calibrate for ground reaction forces either focused on gait analysis (Collins et al. 2009), where shear force magnitudes are expected to be larger, or used center-of-pressure error as the ultimate performance measure and did not quantify uncertainty in ground reaction forces (Cedrado et al. 2009). Furthermore, among the studies that quantified center-of-pressure measurement uncertainty, many relied on point-load application, which is quite dissimilar to the loads that a force plate would experience when a human is standing on it (Bobbert & Schamhardt 1990, Chockalingam et al. 2002, Cedrado et al. 2009, Blanchard Sanhueza 2010, List et al. 2017). There is evidence that pressure distribution affects center-of-pressure errors (Middleton et al. 1999, Schmiedmayer & Kastner 2000), thus more human-like pressure distributions would be informative. More recent studies have employed dynamic loading via a rotating mass or a pendulum to test force plates, but still limited their focus on center-of-pressure measurement (Morasso et al. 2004, Ghersi et al. 2017, Limpach et al. 2025).

In this study, a force-plate performance assessment method is proposed to test the reliability of force plates in measuring the shear force and frequency-domain intersection-point-height during quiet standing. The method was applied to evaluate a portable force plate. The force place measured sagittal-plane shear forces precisely at the lower frequencies such that the errors in the intersection-point-height were less than 10% up to 2 Hz, enough to capture differences observed in older versus younger adults, and survivors of stroke and Parkinson’s disease compared to age-matched controls (Shiozawa et al. 2024a, Sreenivasan et al. 2024, Bartloff et al. 2025). However, the portable force plate was unable to measure the frontal-plane shear forces observed during quiet standing with feet side-by-side due to a small signal-to-noise ratio. Thus, the device background noise critically limited the reliable frequency range of shear-force measurement. These findings highlight the importance of evaluating force-plate uncertainty prior to using the device, especially for low-amplitude measurements such as quiet standing.

## 2 Proposed Method

This section outlines the proposed method for testing force plates intended for human standing studies. It involves placing a rigid weighted frame on the force plate and perturbing the frame with horizontal input forces to induce human-like shear forces and center-of-pressure displacements at the surface of the force plate. The force plate shear-force and center-of-pressure measurements are then compared to the actual values to evaluate the measurement errors. Section 2.1 describes the hardware, followed by Section 2.2, which outlines the procedure for characterizing the test-apparatus dynamics; finally, Section 2.3 presents the procedure for testing force plates using this apparatus.

### 2.1 Setup

Figure 1 provides a schematic of the test apparatus, which comprised a rigid weighted frame with feet to mimic human weight and pressure distribution, a load cell mounted on the frame to measure input force perturbations, and a second frame with a linear actuator that applied horizontal force perturbations via a cable. The weight of the rigid weighted frame and the dimensions of the feet mimicked the human demographic of interest.

**Figure 1.**
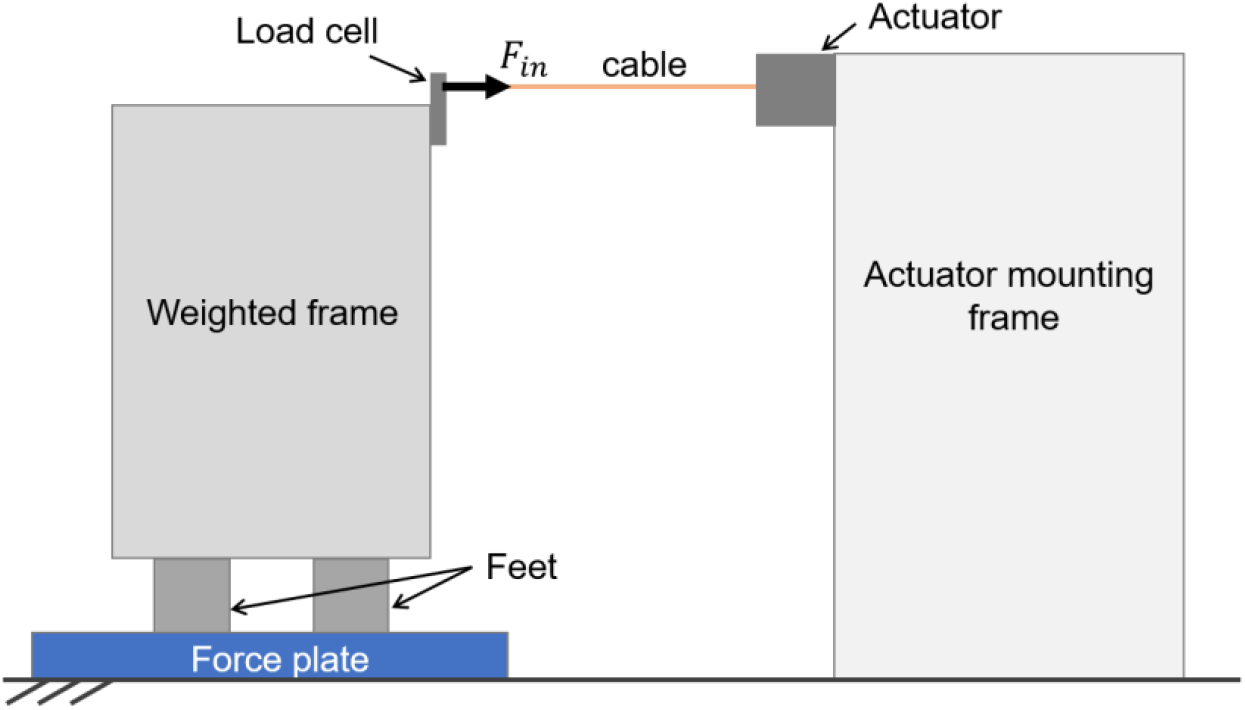
Schematic of test apparatus. A rigid weighted frame was placed on the force plate to be tested. The frame consisted of two feet that mimicked human-like pressure distribution at the force-plate surface, and a load cell that measured input force. A second frame held a linear actuator that applied horizontal force perturbations to the weighted frame via a cable, thereby inducing shear forces and center-of-pressure displacements at the force-plate surface.

From the diagram depicted in Figure 2, the shear force _*Fx*_ and center-of-pressure displacements *x*_*CoP*_ at the force-plate surface are related to the input horizontal forces *F*_*i*_*n* by:

**Figure 2.**
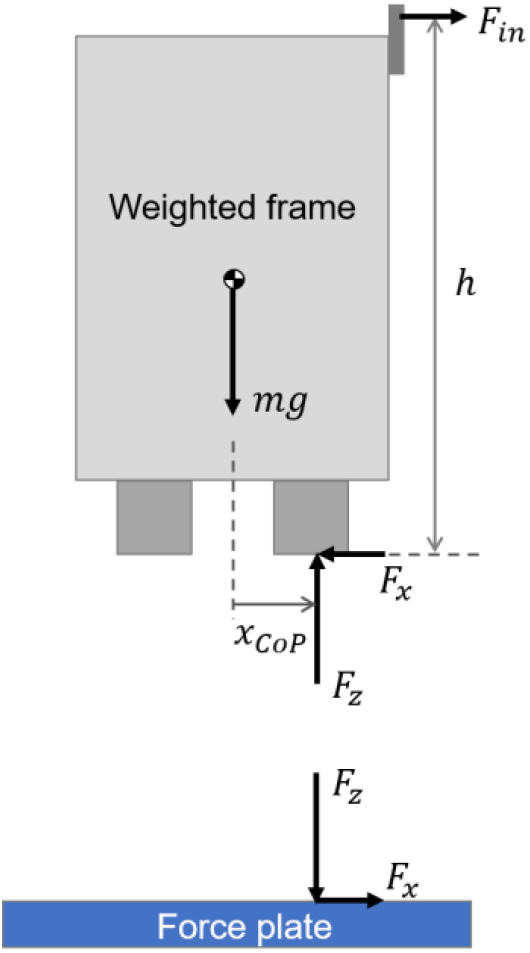
Free-body diagram of the weighted frame and the force plate. *m*: mass of frame *g*: acceleration due to gravity *h*: height of input force from the force-plate surface *F*_*in*_: input force applied via the cable *F*_*x*_: net shear force at the force-plate surface *FZ*: net vertical force at force-plate surface *x*_*CoP*_: center-of-pressure displacement at force-plate surface

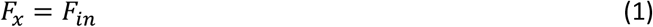

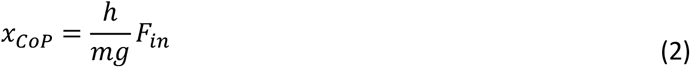

where *m* is the mass of the weighted frame, *g* = 9.81 m/s^2^, and *h* is the height of the input force from the force-plate surface. The input force F_*i*_*n* was limited to 10 N considering the load-cell and actuator capacity. To induce center-of-pressure displacements of up to 1 cm (magnitudes comparable to those observed during human standing) on a 70-kg frame, the height of the force application point had to be around *h* ≈ 0.7 m above the force-plate surface.

### 2.2 Characterization of test apparatus dynamics

Under static conditions, equations (1) and (2) give good approximations to the induced shear force and center-of-mass displacement given the input force, which was measured by the load cell. The load-cell precision, accuracy, hysteresis, and repeatability were characterized via static loading with known masses ranging from 1 g to 1 kg. A load-cell precision of 0.025 N was acceptable for this application.

The objective of the proposed force-plate test was to evaluate measurement uncertainties under both static and dynamic loading. While equations (1) and (2) give good approximations for the induced shear force and center-of-mass displacement under static conditions, the dynamics of the test apparatus must be characterized over the frequency range of interest (up to 8 Hz; Boehm et al. 2019) to estimate the induced shear force and center-of-mass displacement under dynamic conditions. This was accomplished through a sine-sweep test with input sinusoidal forcing of varying frequency applied by the linear actuator via the cable, and the output shear force and center-of-pressure displacement at the bottom of the frame measured using a force plate. To determine whether the dynamics observed were predominantly those of the test apparatus or of the force plate, two different force plates were tested. Note that since the cable allowed forces to be applied in only one direction, input sine-wave forcing was applied with a DC offset; this will not affect the results as long as the system is sufficiently linear. The sine-sweep test was repeated along the two axes of the force plates with the test apparatus oriented in two different directions (feet parallel to the forcing direction versus feet perpendicular to the forcing direction) to assess directional differences.

### 2.3 Evaluation of force-plate measurement

Two different tests evaluated the force-plate uncertainties under quasi-static and dynamic loading conditions.

To quantify uncertainty under quasi-static loading conditions, a ramp-and-hold test was used. We applied a gradual ramp and hold for at least 10 seconds at several different ramp end-point amplitudes to test the measurement accuracy under varying force amplitudes. The force-plate shear-force and center-of-pressure measurement errors were quantified via a root-mean-squared error relative to the estimated true values as given by equations (1) and (2) and the load-cell measurements.

To test the force plates under human-like loading conditions, a dynamic-loading test with input forces designed to produce shear forces and center-of-pressure displacements mimicking human standing was used. Example human shear-force and center-of-pressure trajectories during quiet standing can be extracted from public data sets such as those of Santos & Duarte (2016) and Santos et al. (2017). The input force required to reproduce the human shear forces or center-of-pressure displacements was calculated based on equations (1) and (2), i.e.:

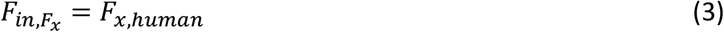

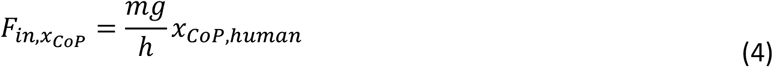

where *Fx,h*_*uman*_ and *x*_*CoP*_,*h*_*uman*_ are human data. Note that a given input forcing can only reproduce either the shear force or the center-of-pressure displacement, and not both simultaneously. Thus, the accuracy of shear-force measurements and center-of-pressure measurements were tested separately.The force-plate measurement errors under dynamic loading were quantified by comparing the power spectral densities of the measurements versus their estimated true values, and computing the error over frequency. The estimated true values were computed based on the load-cell measurements while accounting for any significant dynamics observed in the frequency range of interest. We evaluated errors in the frequency domain because the intersection-point-height (the balance measure of interest) is analyzed in the frequency domain, and error over frequency provides more insight into how measurement uncertainty affects this balance measure.

## 3 Case Study

In this section we describe a test apparatus built to evaluate a portable force plate for its performance in measuring quiet standing behavior. We were particularly interested in whether the device produced reliable measurement of the intersection-point-height, a balance measure that can differentiate between younger and older adults, as well as survivors of stroke and Parkinson’s disease, compared to age-matched able-bodied adults (Shiozawa et al. 2024a, Sreenivasan et al. 2024, Bartloff et al. 2025, Shiozawa et al. 2026). In this work, a modified intersection-point-height measure defined by Shiozawa et al. (2026) was used (see Appendix A for details). Given that the intersection-point-height measure is a quantity derived from shear force and center-of-pressure displacement, we investigated the uncertainty in the raw measurements of shear force and center-of-pressure displacement as well.

### 3.1 Setup

The two frames of the test apparatus were built out of T-slotted framing material (Figure 3). A thick aluminum plate was attached to the base of the weighted frame to provide attachment points for the feet, which were made of 10-cm-wide and 25-cm-long T-slotted framing material. The frame was weighed with five 25 lbs weights to bring the total weight of the structure to 680 N. The weighted frame was instrumented with a uniaxial load cell (ALAMSCN 1kg capacity) installed such that the force-application point was at a height of 0.735 m from the bottom of the feet. A voice coil actuator (SUPT Motion VCAR0055-0249-00A) with 12 N continuous force rating was mounted on the second frame at an adjustable height; once the weighted frame was placed atop the force plate, the height of the actuator was adjusted to make sure that the cable connecting the load cell to the actuator was level. A stainless-steel nylon-coated wire rope (0.044” diameter) was used for the cable. An Arduino Mega microcontroller was used both for actuator control and load-cell data acquisition.

**Figure 3.**
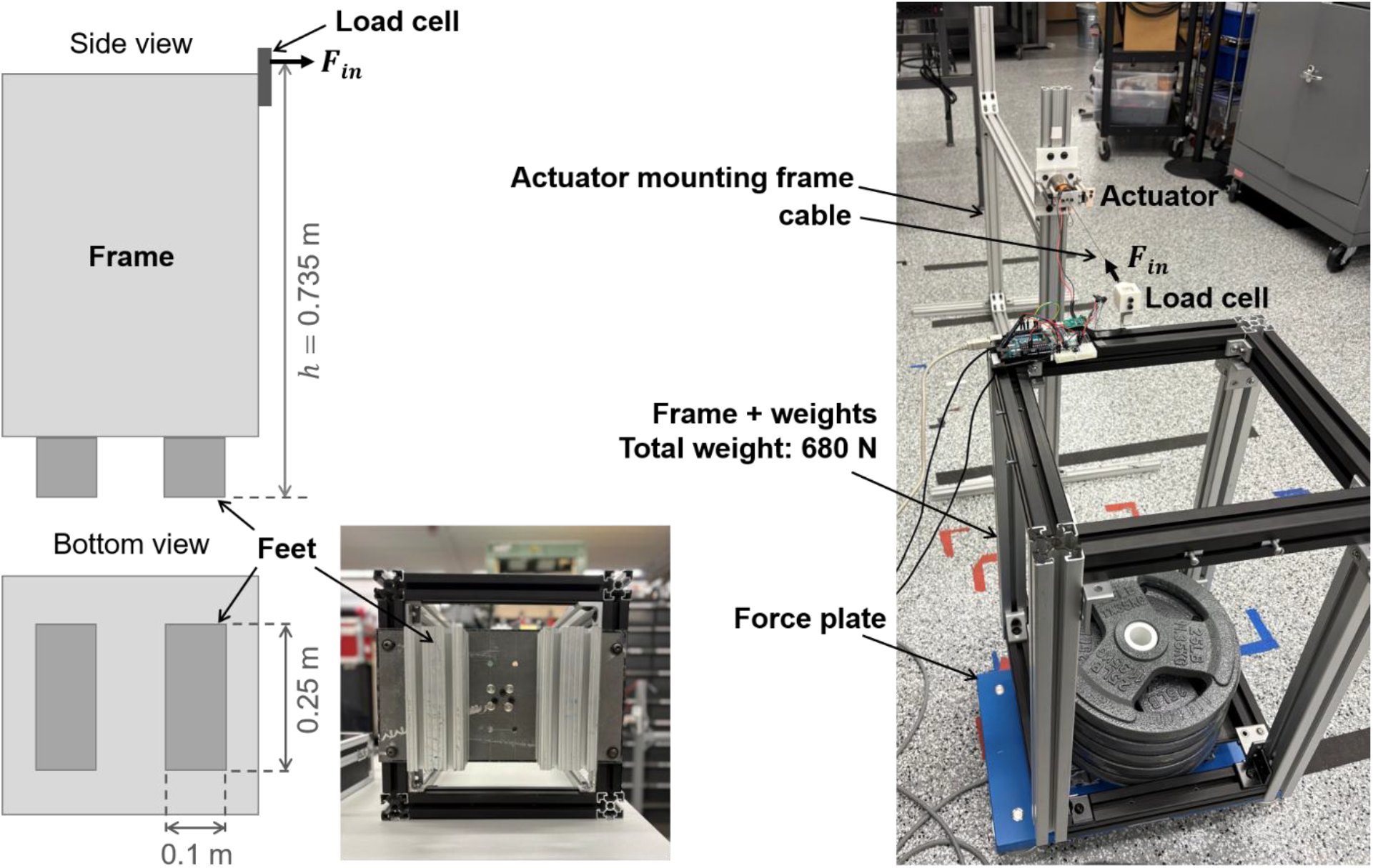
Test apparatus setup. Left: diagram of the weighted frame showing the critical dimensions: the height of the force application point, which dictates the center-of-pressure displacements relative to the input force, and the size of the feet, which determines the pressure distribution at the force-plate surface. Right: photo of the actual test apparatus on a portable force plate (Kistler 9260AA3).

### 3.2 Test apparatus dynamics

First, the load cell was tested via static loading with known masses ranging from 1g to 1kg to evaluate the precision and accuracy of the sensor. Raw data sampled at 80 Hz exhibited errors below 0.025 N, which were further reduced via filtering. The load cell exhibited negligible hysteresis and excellent repeatability across days. Further details of load cell testing can be found in Appendix B.

Thereafter, dynamic testing of the test apparatus was performed through sinusoidal perturbations with an amplitude of 3 N at 12 frequencies (0.01 Hz, 0.04 Hz, 0.1 Hz, 0.4 Hz, 1 Hz, 2 Hz, 3 Hz, …, 8 Hz) applied with a DC offset to account for the unidirectional cable force. The sine-sweep test was performed on two different force plates (Kistler 9260AA3 and Bertec FP4060-07-1000) to measure the shear force *F*_*x*_ and center-of-pressure displacements *x*_*CoP*_ produced at the test-apparatus-force-plate interface. The force-plate measurements *F*_*x*,measured_, *x*_*CoP*_,measured were compared against the load-cell-derived quantities*F*_*i*_nand 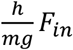 to evaluate the dynamics of the combined test-apparatus-and-force-plate system in the frequency range tested.

The sine-sweep test revealed that the magnitudes of *Fx*,measured and *x*_*CoP*_,measured increased with frequency from 2 Hz onwards, suggesting resonant behavior. Figure 4 shows the scale factors by which the measured quantities *Fx*,measured, *x*_*CoP*_,measured differed from the load-cell-derived quantities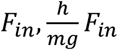 The scale factors were determined by time-aligning the force-plate and load-cell-derived data and scaling the load-cell-derived data to minimize the root-mean-squared error between the two signals. The phase difference between the force-plate shear-force and center-of-pressure measurements revealed no significant difference.

**Figure 4.**
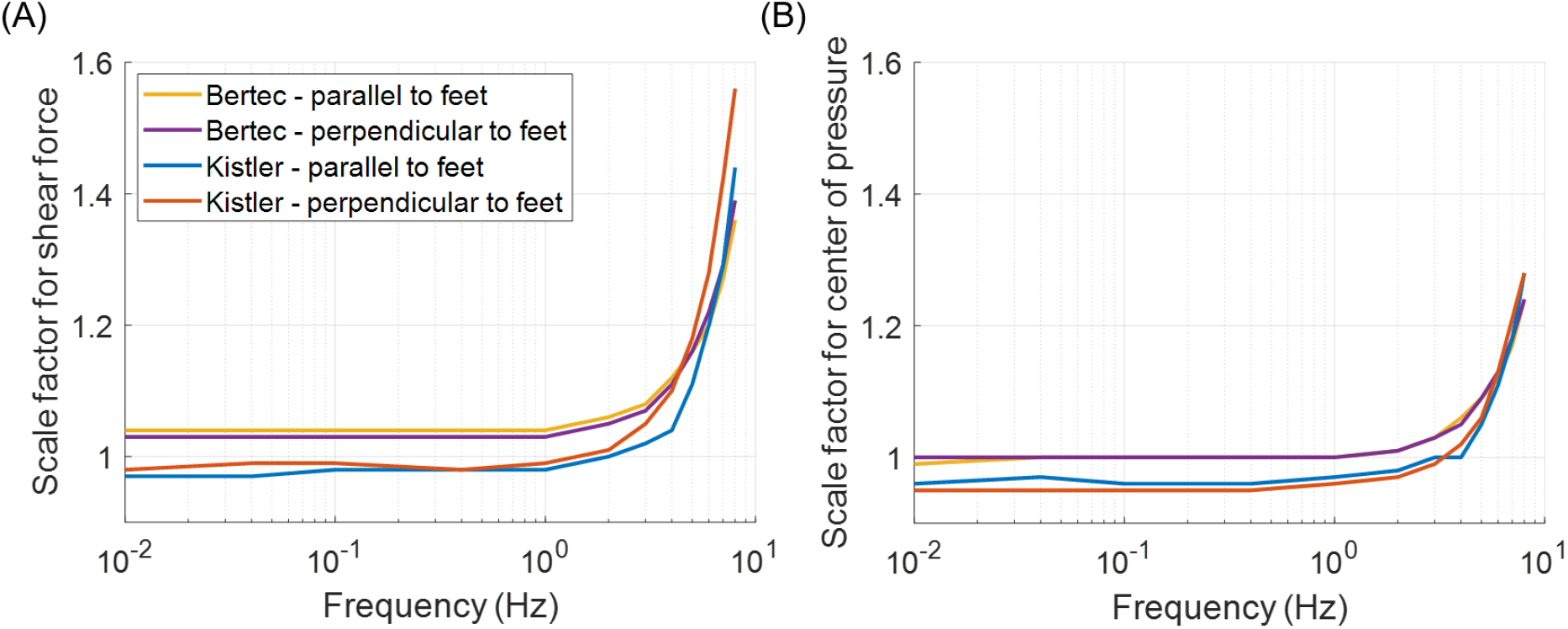
Scale factors by which the measured shear force *F*_*x,measured*_ and -of-pressuredisplacement *x*_*CoP*_,*measured* differed from the load-cell-derived quantities *F*_*i*_*n* and 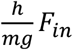Yellow: input force along Bertec force plate y-axis, parallel to testbed feet. Purple: input force along Bertec force plate x-axis, perpendicular to testbed feet. Blue: input force along Kistler force plate x-axis, parallel to testbed feet.Red: input force along Kistler force plate y-axis, perpendicular to testbed feet.

The 3-N excitation used in the sine sweep was much larger than the noise levels of the two force plates, and both force plates from different manufacturers showed consistent responses. Therefore, the observed dynamics were attributed to the test apparatus itself, rather than measurement noise or force-plate characteristics. Furthermore, given the level of agreement in the measurements in the two different directions, we determined that the dynamics were independent of the direction of forcing.

### 3.3 Force-plate measurement errors

To assess the clinical applicability of portable force plates to diagnose balance impairments, we tested a portable device (Kistler 9260AA3) using the method above. A floor-mounted device (Bertec FP4060-07-1000) was also tested to provide a baseline comparison.

The quasi-static ramp-and-hold test was performed at four different amplitudes: 0 N, 1 N, 4 N, 8 N. The force-plate shear-force and center-of-pressure data were aligned with the load-cell-derived quantities to compute the root-mean-squared error in the force-plate measurements. The ramp-and-hold test was repeated in two different forcing directions (parallel versus perpendicular to feet) for the full weighted frame (680 N) and a lightened frame (140 N) with the 25-lbs weights removed, to test the effect of vertical load on force-plate measurement accuracy. Tables 1 and 2 summarize the results. The root-mean-squared errors were also computed for a low frequency (0.01 Hz) sinusoidal input force for comparison. The portable device showed larger errors in both shear force and center-of-pressure displacements compared to the floor-mounted device. While shear force errors were relatively consistent across the three conditions, the center-of-pressure displacement errors were larger for the light-weight frame, consistent with prior work (Chockalingam et al. 2002).

**Table 1.** Root-mean-squared error (RMSE) in shear-force data from each force plate in the two directions.

| Shear force RMSE (N) | Portable device |  | Floor-mounted device |  |
| --- | --- | --- | --- | --- |
|  | Direction 1 (parallel to feet) | Direction 2 (perpendicular to feet) | Direction 1 (parallel to feet) | Direction 2 (perpendicular to feet) |
| Ramp + hold (light) | 0.19 | 0.15 | 0.096 | 0.096 |
| Ramp + hold (heavy) | 0.18 | 0.18 | 0.099 | 0.090 |
| 0.01 Hz sinusoid (heavy) | 0.18 | 0.20 | 0.10 | 0.086 |

**Table 2.** Root-mean-squared error (RMSE) in center-of-pressure displacement data from each force plate in the two directions.

| Center-of-pressure displacement RMSE (mm) | Portable device |  | Floor-mounted device |  |
| --- | --- | --- | --- | --- |
|  | Direction 1 (parallel to feet) | Direction 2 (perpendicular to feet) | Direction 1 (parallel to feet) | Direction 2 (perpendicular to feet) |
| Ramp + hold (light) | 0.73 | 0.87 | 0.25 | 0.36 |
| Ramp + hold (heavy) | 0.14 | 0.16 | 0.078 | 0.038 |
| 0.01 Hz sinusoid (heavy) | 0.13 | 0.14 | 0.045 | 0.040 |

For the dynamic-loading test, data from a young healthy male subject standing with eyes open and feet side-by-side for three trials of 60 seconds each were used as the reference human data (Subject 1 from Santos & Duarte 2016). The human shear-force data were directly used as the input force with a DC offset to account for the one-directional cable forcing. The human center-of-pressure displacement data were converted to the input shear force via equation (2), and then high-pass filtered with a 0.4 Hz cut-off to keep the input force signal range within the hardware limits. Input force derived from sagittal-plane data was applied along the direction parallel to the test-apparatus feet, and those derived from frontal-plane data were applied perpendicular to the feet.

Figure 5 compares the average power spectral densities (PSDs) of the force-plate measurements against the estimated true values. The true PSDs were derived by first computing the PSDs of the quantities derived from equations (1) and (2), which were then scaled by the square of the mean scale factors from Figure 4 to correct for the test-apparatus dynamics. The center-of-pressure measurements exhibited smaller errors for a larger bandwidth than shear-force measurements. Errors were more pronounced at higher frequencies for both quantities, and the PSDs showed a clear noise floor. Frontal-plane shear-force and center-of-pressure data showed lower power and a similar noise floor, resulting in larger errors compared to sagittal-plane data.

**Figure 5.**
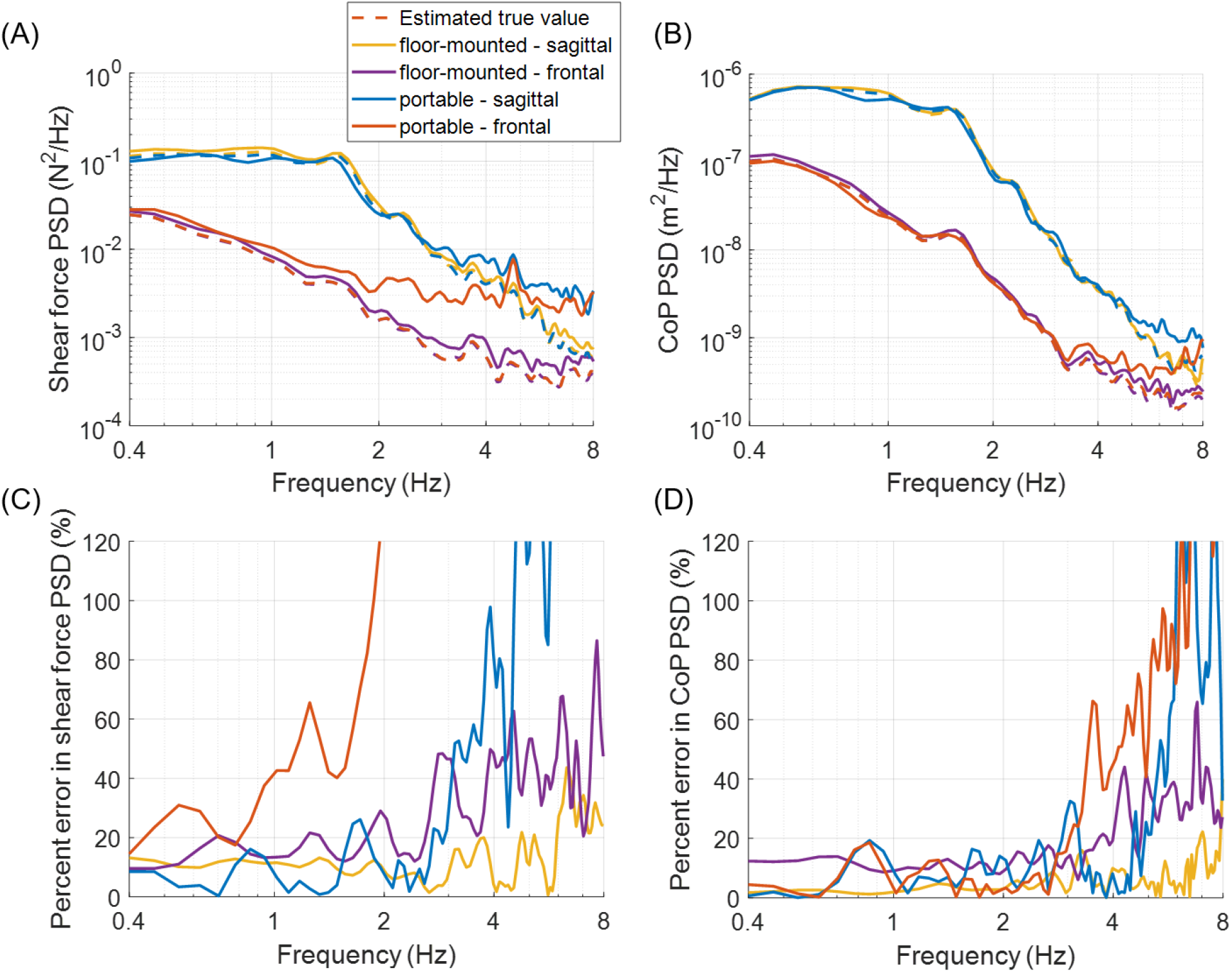
Shear force and center-of-pressure (CoP) displacement power spectral density (PSD) from force-plate data (solid lines) versus estimated true values (dashed lines) when input force was applied to mimic shear forces and CoP displacements seen during human standing. A and B show the PSDs, while C and D show the percent errors between the force-plate data versus the estimated true values. Solid yellow: input force derived from sagittal-plane data applied atop floor-mounted force plate.Solid purple: input force derived from frontal-plane data applied atop floor-mounted force plate. Solid blue: input force derived from sagittal-plane data applied atop portable force plate.Solid red: input force derived from frontal-plane data applied atop portable force plate.Dashed: estimated true values derived from load-cell data.

Force-plate data generated from the trials mimicking human-center-of-pressure were subsequently low-pass-filtered to 10 Hz and used to compute ten statistical CoP-based measures from Prieto et al. (1996), while the force-plate data generated from the trials mimicking human-shear-force were used to compute the frequency-domain intersection-point-height measure. Figure 6 presents the errors in the force-plate-derived measures compared to the estimated true values from the load cell. The center-of-pressure measures that depended more on lower-frequency content, such as mean (mdist) or root-mean-square distance (rdist), exhibited low errors (<5%); measures sensitive to higher-frequency signals like mean velocity (mvelo) exhibited larger errors (for the portable device, up to 27% for sagittal-plane data and almost 60% for frontal-plane data). The portable device produced the intersection-point-height measure with errors less than 10% below 2 Hz for the sagittal-plane data, but was unable to produce reliable measures from the frontal-plane data. The portable device generally exhibited larger errors than the floor-mounted device.

**Figure 6.**
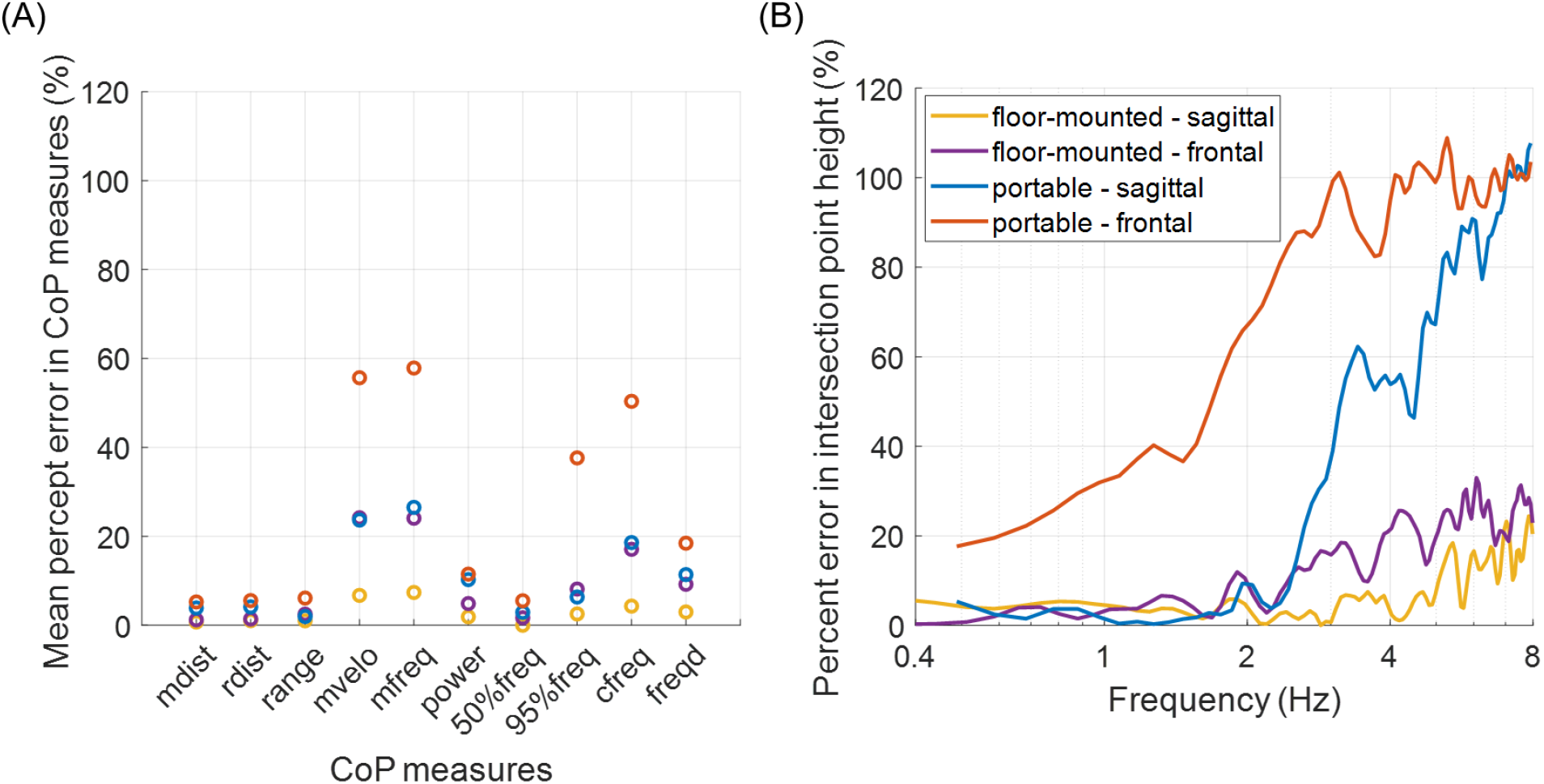
Percent error in center-of-pressure (CoP) measures and intersection-point-height derived from force-plate data versus estimated true values derived from load-cell data.Blue: input force derived from sagittal-plane data applied atop floor-mounted force plate. Red: input force derived from frontal-plane data applied atop floor-mounted force plate. Yellow: input force derived from sagittal-plane data applied atop portable force plate.Purple: input force derived from frontal-plane data applied atop portable force plate. Center-of-pressure measures from Prieto et al. 1996:mdist: mean distance rdist: root-mean-squared distance range: difference between maximum and minimum distances mvelo: mean velocity mfreq: mean frequency power: total power50%freq: 50%-power frequency 95%freq: 95%-power frequency cfreq: centroidal frequency freqd: frequency dispersion

## 4 Discussion

Shear-force data provides useful information about balance behavior in the form of the intersection-point-height measure. Low-cost portable force plates with reliable shear-force measurements would enable regular balance check-ups to detect balance impairments before a fall. The case study results above revealed that existing portable force plates may be at a critical point: the portable device tested here produced a reliable intersection-point-height measure up to 2 Hz for sagittal-plane data, yet was unable to produce an intersection-point-height with acceptable accuracy for frontal-plane data. The power spectral densities showed a clear noise floor at higher frequencies, suggesting that background noise might be the reason for the larger errors at higher frequencies. Indeed, by adding white noise with the same signal strength as the devices’ background noise level to the original clean signals, we were able to reproduce the errors observed in the intersection-point-height measure from both the portable and the floor-mounted devices (Figure 7). This suggests that, although force-plate measurement error may result from several different factors, such as nonlinearities, hysteresis, and crosstalk, the noise floor is the limiting factor for standing applications. The observed broadband noise that was present even with no load on the force plate likely comes from the device electronics, and may benefit from improved shielding or other adjustments. The significant measurement noise also explains the inability to precisely measure the frontal-plane intersection-point-height with the portable device: the shear forces observed in the frontal plane during human quiet standing had a much smaller variance, resulting in a signal-to-noise ratio that was too small to reliably measure with the force plate. In these experiments, data were taken from a subject standing with eyes open and feet side-by-side with heels 10 cm apart. The eyes-closed data in the same dataset did not show significant increases in shear-force magnitude. Measuring balance behavior in tandem or single-leg stance induced greater sway, thereby increasing the signal-to-noise ratio (Shiozawa et al. 2024b).

**Figure 7.**
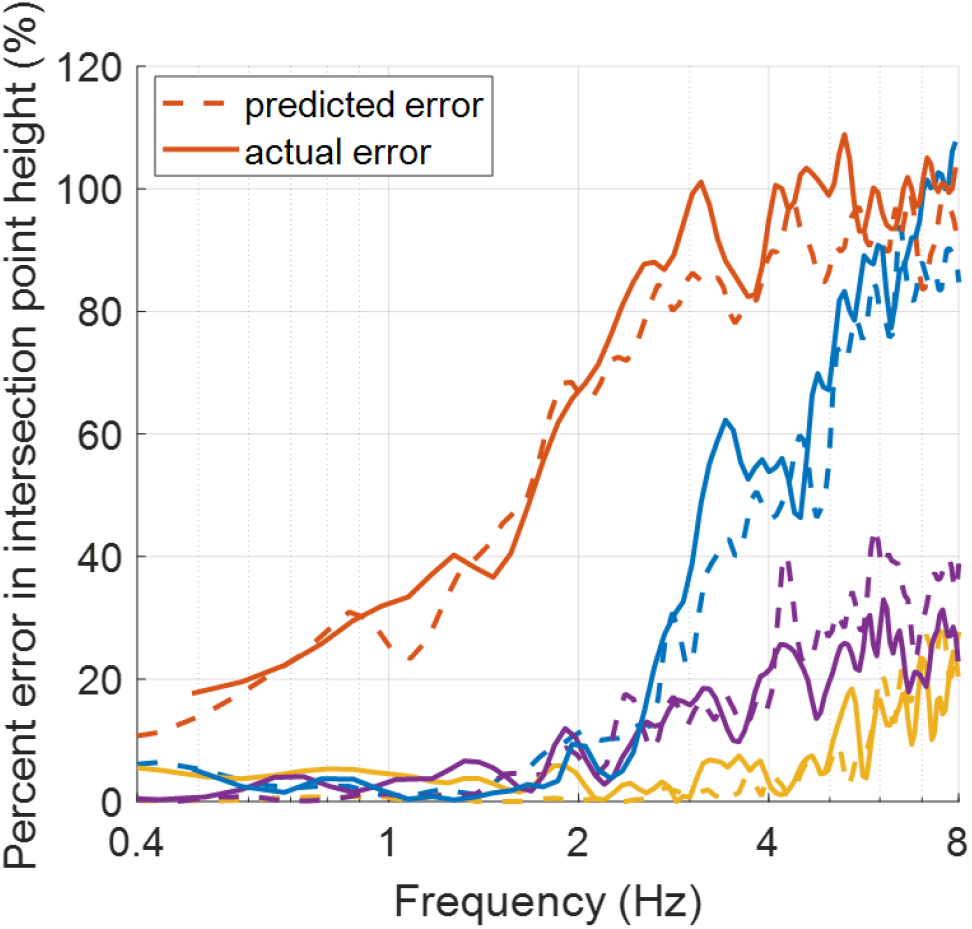
Percent error in intersection-point-height predicted using artificial data generated from the load-cell-derived true values with additive white noise of the same variance as the noise floor observed in the force plates (dashed) compared to the percent errors from force-plate data (solid). Yellow: input force derived from sagittal-plane data applied atop floor-mounted force plate.Purple: input force derived from frontal-plane data applied atop floor-mounted force plate. Blue: input force derived from sagittal-plane data applied atop portable force plate.Red: input force derived from frontal-plane data applied atop portable force plate.

The force-plate performance assessment revealed the limitations of an existing portable force plate, thus highlighting the need to assess whether a given device is appropriate for standing applications. The method presented here provides a means to characterize the force-plate measurement uncertainties for quasi-static and human-like dynamic loading. The method was able to characterize the force-plate errors in both shear force and center-of-pressure measurements, and identified the noise floor as the dominant source of the measurement errors. The method can also be applied to directly estimate uncertainties of commonly used balance measures to evaluate how reliable each measure is when obtained using a specific force plate.

One limitation of the method is that the test-apparatus dynamics must first be characterized using a reliable force plate. Ideally, the sine-sweep test should be repeated on multiple different force plates to isolate the test-apparatus dynamics from possible force-plate dynamics. Given that the estimation of the true shear force and center-of-pressure displacement at the force-plate surface depends on the identified dynamics of the test apparatus, careful characterization is important. Nevertheless, if the test apparatus built is rigid enough, its dynamics will have minimal effect on the result, and equations (1) and (2) provide acceptable estimates of the true values. These can be used to provide conservative estimates of the force-plate measurement errors independent of the accuracy of the test-apparatus-dynamic characterization. In the case study presented here, using equations (1) and (2) to estimate the true values without correcting for the test-apparatus dynamics produced similar results for the force-plate measurement errors. It is plausible that the human-like input force signal was attenuated sufficiently at the higher frequencies such that any resonant behavior of the weighted frame did not induce significant amplification of the output shear force.

Another limitation of the method is the fact that the dynamic loading used is not entirely human-like, as a DC offset must be included to apply forces via a cable, and the shear force and center-of-pressure displacements observed in human standing cannot be reproduced simultaneously. Despite this limitation, the dominant source of measurement error, the noise floor, was identified and reproduced numerically.

## 5 Conclusion

Falls have dire consequences for one’s life. As the global population ages, the number of fall-related injuries and deaths will only rise. Early detection of balance impairments will allow for preventive measures. Force plates have a potential to be used as a diagnostic tool for balance disorders by measuring intersection-point-height, which can detect differences in balance behavior of healthy versus impaired persons. Portable force plates are convenient for their simple ‘plug-and-play’ capability without the need for permanent installation in the floor. The analysis presented here revealed that a commonly-used laboratory-grade portable force plate was able to measure sagittal-plane intersection-point-height below 2 Hz with acceptable accuracy despite the small shear-force magnitudes observed during quiet standing. However, the device could not measure shear forces in the frontal plane accurately due to the low signal-to-noise ratio. Future studies should evaluate the performance of other existing portable force plates and investigate cost-effective designs that achieve more precise shear-force measurements during quiet standing.

## Acknowledgements

The authors would like to thank Evan Linton for his assistance in data collection, and the MIT Center for Clinical and Translational Research and the MIT Biomimetic Robotics Laboratory for lending us their force plates. This work was partially supported by the Eric P. and Evelyn E. Newman Fund, the Bernard (Ben) Gold Fellowship, and the MathWorks Engineering Fellowship.

## Appendix A intersection-point-height

The intersection-point-height measure is a measure of quiet-standing balance developed by Boehm et al. given by the slope of the best-fit line through band-pass-filtered center-of-pressure displacement and foot-force orientation measured by the angle of the force vector with respect to the vertical; by using a band-pass filter centered at different frequencies, the intersection-point-height value can be computed over frequency (Boehm et al. 2019). The original intersection-point-height was based on the slope of the best-fit-line defined by principal component analysis; i.e., one that minimized the sum of orthogonal distances from all data points to the line (Boehm et al. 2019). Here, we use the modified intersection-point-height used in Shiozawa et al. (2026), where the best-fit-line is defined by the linear regression line; i.e., the line that minimized the vertical distances from all data points to the line. The slope of the regression line of the center-of-pressure-displacement versus foot-force-orientation data is equivalent to the ratio of the covariance between center-of-pressure displacement and foot-force orientation divided by the variance of the foot-force orientation. Alternatively, instead of band-pass-filtering the data at each frequency, the intersection-point-height over frequency *Z*_*IP*_(*f*) can be obtained through spectral analysis (Sugimoto-Dimitrova et al. 2024). The modified intersection-point-height can be computed from the cross-spectral density between center-of-pressure displacement and foot-force orientation *G*_*xCoP*θ_^*F*^ (*f*) and the auto-spectral density of foot-force orientation *G*θ^*F*^θ^*F*^ (*f*) as:

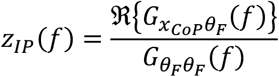

where ℜ{*G*_*x*^*CoP*^_θ^*F*^ (*f*)} is the real part of the complex quantity *G*_*x*^*CoP*^_θ^*F*^ (*f*), and *x*_*CoP*_ and θ_*F*_ denote the center-of-pressure displacement and foot-force orientation, respectively. The foot-force orientation is related to shear component of the foot-force *Fx* by:

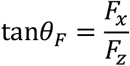

*F*_*Z*_where *F*_*Z*_ denotes the vertical loads. Given that the vertical loads are much larger than the horizontal loads during quiet standing, the foot-force orientation can be approximated as θ_*F*_ ≈ *Fx*/*F*_*Z*_.

## Appendix B load-cell testing

The uniaxial load cell (ALAMSCN 1kg capacity) was calibrated using static loading with known weights ranging from 1 g to 1 kg. A calibration factor was computed that minimized the error between measured and true loads. Figure B.1 shows the distribution of load-cell measurement errors for the calibrated load cell. The raw load-cell measurements sampled at 80 Hz exhibited errors below +/-0.025N in the negative loading direction, while in the positive loading direction errors up to +/-0.05 N were observed. The load cell was mounted on the weighted frame such that the cable pulled in the negative direction to ensure smaller errors.

**Figure B.1.**
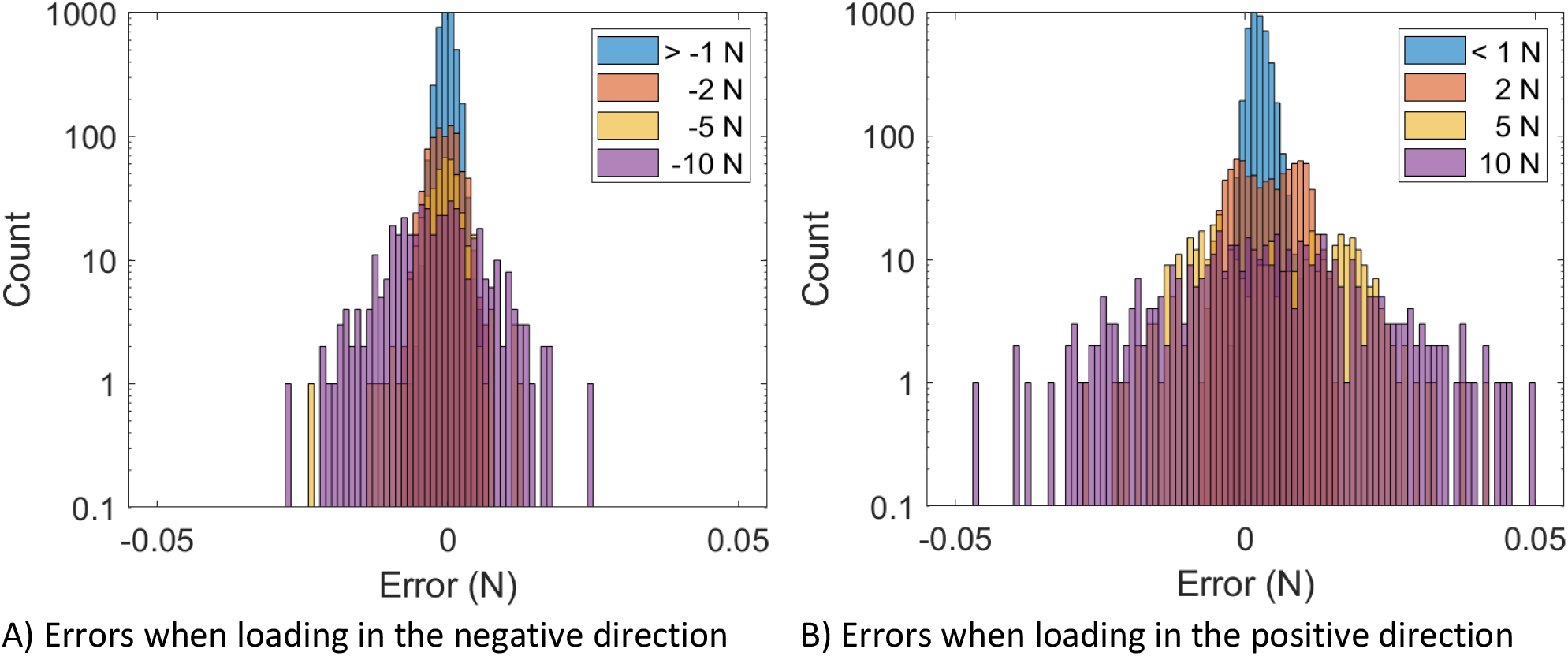
Distribution of load-cell-data errors during static-loading tests when weights were loaded in two different directions: A) in the “negative” loading direction of the load cell, and B) in the “positive” loading direction of the load cell. Legend indicates the load.

With the calibration factor determined, the load cell was then tested via static loading of the same known weights to test for repeatability and hysteresis.

The static-loading test was repeated four times over the course of a month. The load cell exhibited excellent repeatability, showing negligible variation in measurement error across trials.

To test hysteresis, the weights were placed on the load cell one after another cumulatively, from smallest weights to largest, and then unloaded from largest to smallest. The loading and unloading were repeated twice. Weights were added/removed at 10-second intervals, and the load-cell measurements were averaged over the 10 seconds to obtain a measurement for each weight variation. Figure B.2 shows the load-cell measurements plotted against the true load, where minimal differences were observed during loading and unloading, indicating negligible hysteresis.

**Figure B.2.**
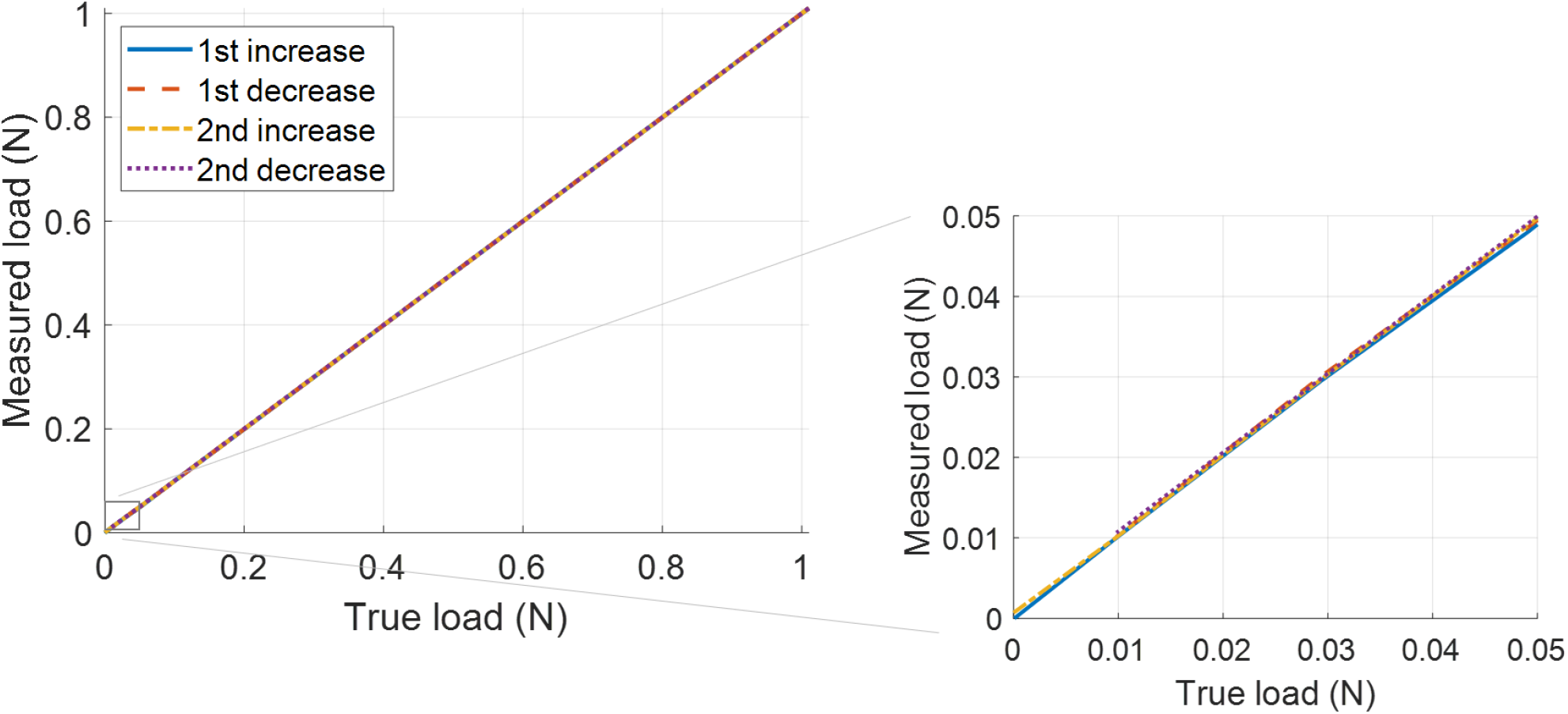
Load cell hysteresis test. The measured loads during the loading (increase) and unloading (decrease) of weights are plotted against the true loads, showing minimal difference in load-cell measurements when loads were increased versus decreased. The right plot shows a zoomed-in plot for loads below 0.05 N.

